# Rapid mosaic brain evolution under artificial selection for relative telencephalon size in the guppy (*Poecilia reticulata*)

**DOI:** 10.1101/2021.03.31.437806

**Authors:** Stephanie Fong, Björn Rogell, Mirjam Amcoff, Alexander Kotrschal, Wouter van der Bijl, Séverine D. Buechel, Niclas Kolm

**Author notes:** Corresponding author: Stephanie Fong.

## Abstract

The vertebrate brain displays enormous morphological variation and the quest to understand the evolutionary causes and consequences of this variation has spurred much research. The mosaic brain evolution hypothesis, stating that brain regions can evolve relatively independently, is an important idea in this research field. Here we provide experimental support for this hypothesis through an artificial selection experiment in the guppy (*Poecilia reticulata*). After four generations of selection on relative telencephalon volume (relative to brain size) in replicated up-selected, down-selected and control-lines, we found substantial changes in telencephalon size, but no changes in other regions. Comparisons revealed that up-selected lines had larger telencephalon while down-selected lines had smaller telencephalon than wild Trinidadian populations. No cost of increasing telencephalon size was detected in offspring production. Our results support that independent evolutionary changes in specific brain regions through mosaic brain evolution can be important facilitators of cognitive evolution.

## Introduction

The vertebrate brain contains a number of morphologically and functionally distinct regions, linked through intricate connective systems, ranging from shorter local links between adjacent structures to connections between spatially distant regions through myelinated axons. Although the vertebrate brain is surprisingly conserved with regards to the organization of the regions it contains, there is large variation among vertebrate species in the relative size and function of different regions^1^. One example is the disproportionately large neocortex in large-brained primates such as humans^2^. But how has such variation evolved among the vertebrates and what are the functional consequences of this variation? Despite over a century of research, these remain to be important questions in the study of brain evolution^3^. One of the leading brain evolution theories, the mosaic brain evolution hypothesis^4^, posits that changes in individual regions may occur due to region-specific selection (e.g. according to explicit cognitive or sensory requirements), relatively independent of changes in other regions. If true, this should result in a heterogeneous pattern of brain evolution^4,5^.

The mosaic brain evolution hypothesis has received ample support mainly from comparative analyses at the interspecific level, by means of examining how ecological factors correlate with specific alterations in neural anatomy across species or populations, e.g. ^6,7–11^. One important example of empirical evidence for this hypothesis was provided by Barton and Harvey ^4^, where the authors analyzed the evolutionary changes in brain regions associated with specific functional units in primates and insectivores. The partly autonomous evolution of these functionally linked components with respect to other brain regions and overall brain size provided support for mosaic brain evolution in mammals. Despite a discernable bias towards mammalian studies, support for mosaic brain evolution in various fish and avian species has also been demonstrated. For example, 3D reconstructions of the brains of African mormyrid fishes that use active electro sensing, revealed an enlarged cerebellum in comparison to outgroup species without such a system^12^. Another well-studied example concerns food caching in birds, where the ability to store food for later retrieval has been found to be positively correlated with hippocampus volume in species performing this cognitively demanding behaviour^13–15^. Recently, also intraspecific comparative analyses have complemented such studies and shown positive correlations between song complexity and the brain regions governing singing behaviour in song birds (i.e. HVC and RA)^16^.

Although interspecific and intraspecific correlative comparative analyses form an important tool to investigate evolutionary patterns and generate hypotheses^6,10,17^, experimental evidence is needed to fully understand the independent evolutionary potential of brain regions. Artificial selection experiments on mice and fish have revealed that increases in relative brain size can occur quickly and yield important cognitive benefits (increased associative learning^18–23^, more accurate mate preferences^24,25^, and more effective predator avoidance^26,27^), but also high energetic costs (lower offspring production^19^, reduced innate immune response^28^, and shorter intrinsic life-span^29^). But in line with the mosaic brain hypothesis, relative brain size is a rather crude measure of brain morphology, and evolutionary changes in brain morphology in wild populations are unlikely to target the entire brain^30,31^. Another fitting and important aspect of the mosaic brain hypothesis concerns the costly nature of neural tissue, which should favor evolutionary expansion of only functionally relevant structures thereby reducing unnecessary energy expenditure^10,32^. In order to increase our understanding of brain morphology evolution, and to go a significant step beyond brain size^33^, artificial selection experiments that target region evolution could be instrumental.

Here we provide such an experimental test of the mosaic brain evolution hypothesis through an artificial selection experiment that targets relative telencephalon size (i.e. telencephalon volume in relation to the brain remainder) in guppies (*Poecilia reticulata*). The established role of the telencephalon as a cognitive centre in fish^34–36^ and other vertebrates^37,38^ makes this region an appealing target of artificial selection with highly general implications, especially given the potential for future tests of the implicated costs and cognitive benefits. We chose the guppy as the model for the experiment because it is a live-bearing teleost species with a well-studied, interesting and variable ecology e.g. ^39,40–42^, it has a relatively short generation time^33^, and large populations can be kept in the lab with ease. Specifically, we aimed to increase and decrease the relative size of the telencephalon in relation to the brain remainder while examining for concomitant changes in other aspects of brain morphology. We also investigated sex specific effects. As a measure of potential reproductive costs^43^ associated with changes in telencephalon size, offspring production, one of the best proxies for fitness in the guppy, was quantified during selection. Finally, we investigated how the artificial selection lines compared to wild populations by comparing their relative telencephalon size against a large data-set spanning across 16 Trinidadian guppy populations.

## Results

After four generations of selection for relative telencephalon size, there was an average difference of 10.1 % between the up-selected and down-selected females, and an average difference of 9.5 % in males. There was an overall significant difference between selection treatments for both females and males (estimated differences between up-selected and down-selected lines: ß, presented with 95 % credible intervals (CI), females: ß = 0.046 [0.0094; 0.089], *P_MCMC_* = 0.044; males: ß = 0.022 [0.0052; 0.041], *P_MCMC_* = 0.028) (Figure 1; Table 1). Comparisons to the unselected control-lines revealed non-significant trends for selection to proceed in an asymmetrical manner in females (up-selected vs. control-line: ß = 0.031 [−0.0023; 0.065], *P_MCMC_* = 0.052; down-selected vs. control-line: ß = −0.014 [−0.077; 0.064], *P_MCMC_ =* 0.50) but no such pattern was detected in males (up-selected vs. control-line: ß = 0.030 [−0.017; 0.098], *P_MCMC_* = 0.18; down-selected vs. control-line: ß = −0.0094 [−0.064; 0.050], *P_MCMC_ =* 0.66). The effects of selection were consistent also when controlling for body length (Table 2), suggesting that relative telencephalon size, both in relation to the brain remainder and body size, were affected by the directional selection procedure.

**Figure 1.**
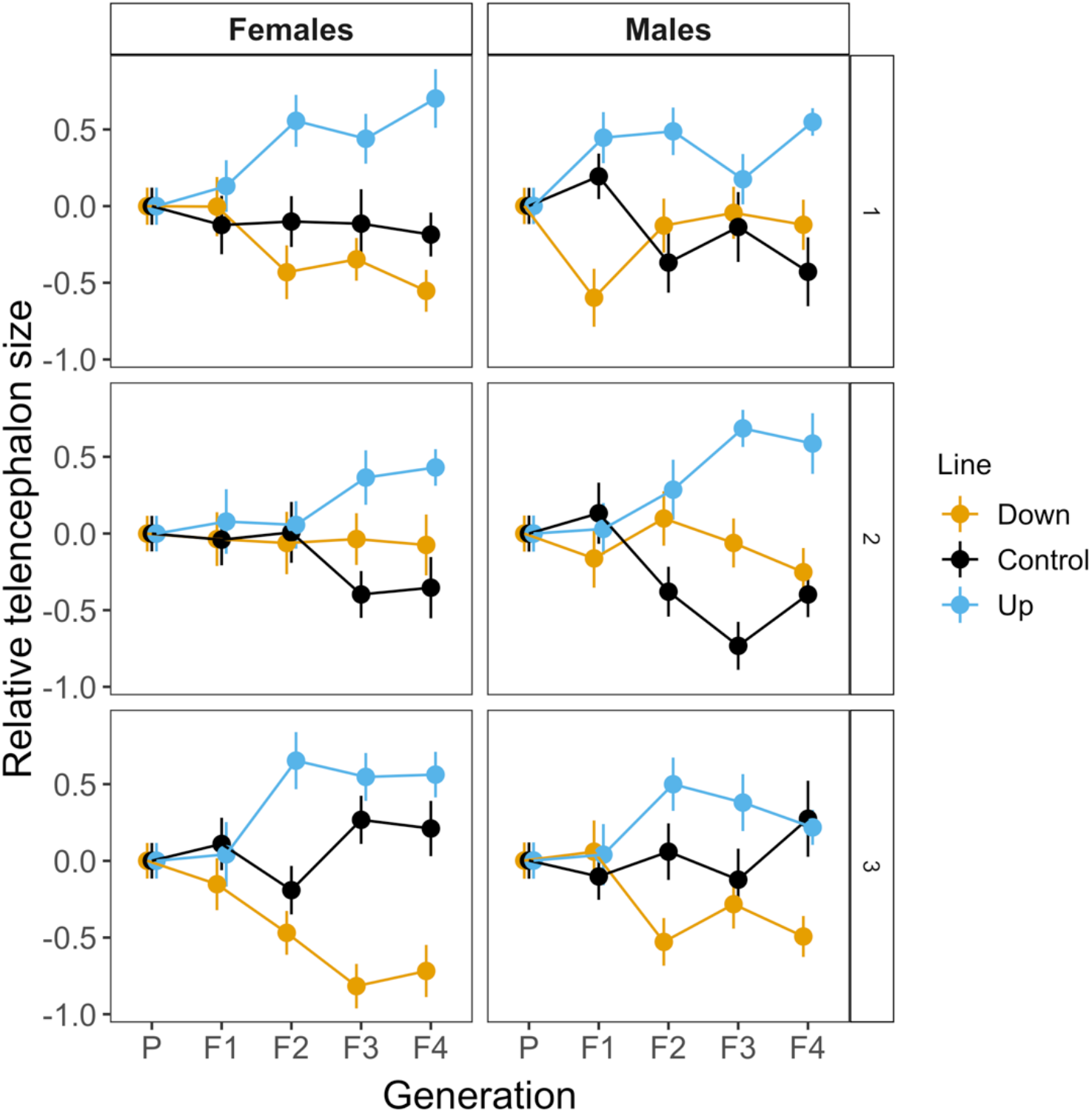
Relative telencephalon size in response to directional selection over four generations. P indicates the starting parental population. Depicted are the mean and standard error values (sem) for standardized residuals of telencephalon volume regressed on the volume of the brain remainder (total brain volume minus telencephalon volume) within each generation and replicate. The figure shows the selection response across the three replicates, with the left and right panel separately illustrating the response in females and males, respectively.

**Table 1.** Model output for relative telencephalon size (relative to the remainder of the brain) in generation F_4_ females and males, presented with mean estimates and 95 % credible intervals (CI). The intercept is set to the intercept of the regression of log telencephalon size on log brain remainder (total brain volume minus telencephalon volume) for the down-selected line in females. Mean estimates are indicative of differences between the intercept (down-selected females) and the specified variable (conforming to the default contrast matrix in R).

| **Adult Generation F<sub>4</sub>** |  |  |  |  |
| --- | --- | --- | --- | --- |
| **Fixed terms** | Estimate ($\beta$) | Lower CI | Upper CI | $P_{MCMC}$ |
| Intercept | -0.703 | -0.901 | -0.556 | 0.004 ** |
| Brain remainder | 0.892 | 0.769 | 1.02 | <0.001 *** |
| Sex Male | -0.0494 | -0.112 | 0.0213 | 0.144 |
| Unselected control-line | 0.0168 | -0.0219 | 0.0551 | 0.272 |
| Up-selected line | 0.0457 | 0.00909 | 0.0848 | 0.030 * |
| Rest of brain:Sex Males | -0.0977 | -0.262 | 0.0624 | 0.282 |
| Sex Males:Unselected control-line | -0.00629 | -0.0253 | 0.0110 | 0.512 |
| Sex Males:Up-selected line | 0.000940 | -0.0172 | 0.0198 | 0.914 |
| **Random terms** | Variance | Lower CI | Upper CI | $P$ |
| Residual | 0.00188 | 0.00166 | 0.00214 | NA |
| Selection treatment:Replicate | 0.000490 | 2.30e<sup>-6</sup> | 0.00160 | NA |
| Replicate | 0.0633 | 4.87e<sup>-12</sup> | 0.155 | NA |

**Table 2.** Model output for relative telencephalon size (relative to body size) in generation F_4_ females and males, presented with mean estimates and 95 % credible intervals (CI). The intercept is set to the intercept of the regression of log telencephalon size on log body size (standard length) for the down-selected line in females. Mean estimates are indicative of differences between the intercept (down-selected females) and the specified variable (conforming to the default contrast matrix in R).

| **Adults Generation F<sub>4</sub>** |  |  |  |  |
| --- | --- | --- | --- | --- |
| **Fixed terms** | Estimate ($\beta$) | Lower CI | Upper CI | $P_{MCMC}$ |
| Intercept | -1.57 | -1.90 | -1.26 | 0.002 ** |
| Standard length | 0.956 | 0.761 | 1.14 | <0.001 *** |
| Sex Males | -0.882 | -1.22 | -0.494 | <0.001 *** |
| Unselected control-line | 0.0149 | -0.0187 | 0.0539 | 0.318 |
| Up-selected line | 0.0571 | 0.0191 | 0.0973 | 0.014 * |
| Standard length:Sex Males | 0.632 | 0.330 | 0.901 | <0.001 *** |
| **Random terms** | Variance | Lower CI | Upper CI | $P$ |
| Residual | 0.00202 | 0.00178 | 0.00228 | NA |
| Selection treatment:Replicate | 0.000675 | 1.21e<sup>-5</sup> | 0.00206 | NA |
| Replicate | 0.0643 | 2.03e<sup>-8</sup> | 0.192 | NA |

Results presented hereafter are based on comparisons between down-selected and up-selected lines only, unless otherwise stated (for a full summary of results, see Table S1, Figure S2 and Figure S3). Overall brain size (controlling for body length) was found to be larger in the up-selected lines (ß = 0.023 [0.0073; 0.042], *P_MCMC_* = 0.022), but further analysis across each sex revealed that this difference in relative brain size was mainly driven by effects in males (males: ß = 0.022 [0.0052; 0.041], *P_MCMC_* = 0.028; females: ß = 0.025 [− 0.0070; 0.058], *P_MCMC_* = 0.098) (Table S2). No significant differences in any of the other brain regions assessed were detected (Table S1).

Realized heritabilities of relative telencephalon size were variable, but congruent between sexes when comparing the up- and down-selected lines against the control-lines (up-selected females: 1.03 [−0.66; 4.35], down-selected females: 0.44 [−1.37; 2.73], up-selected males: 1.42 [−1.78; 7.00], down-selected males: 0.0078 [−1.92; 1.50]).

A previous study comparing the brain morphology of wild-caught and lab-reared guppies found that the latter had smaller telencephalon and optic tectum in relation to the former after just a single generation^44^. Here we compared the mean telencephalon size between selection treatments and 16 Trinidadian wild populations in females and found that the unselected control-lines had a telencephalon size that most closely resembled that found in natural populations, while up-selected and down-selected lines had larger and smaller relative telencephalon size than natural populations (Figure 2). Upon closer examination, it was also found that selection over four generations resulted in a comparable range of relative telencephalon sizes in our selected sample (range −0.76 to 1.3) in comparison to natural populations (range −1.2 to 0.67), contradicting inbreeding issues in this trait over several generations of selection in our experiment (Figure S4).

**Figure 2.**
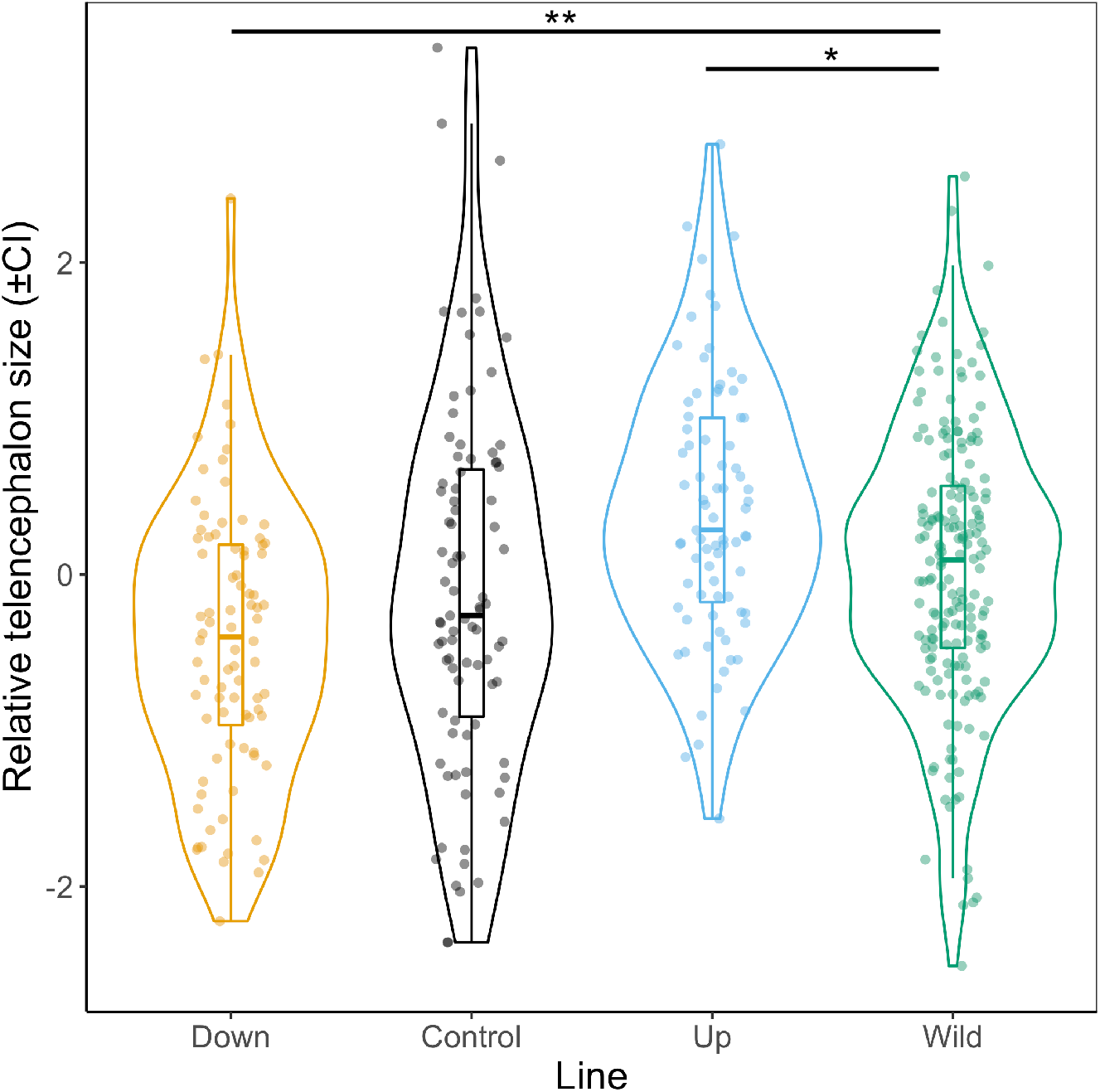
Female relative telencephalon size after three rounds of selection (generation F_4_) in comparison to wild fish obtained from 16 wild populations in Trinidad. Plotted on the y-axis are the estimated marginal means and 95 % confidence intervals, obtained from standardized residuals of telencephalon volume regressed on brain remainder (total brain volume minus telencephalon volume). Pairwise comparisons were performed with respect to the wild populations and *P*-values were calculated following Holm’s adjustment for multiple comparisons. \*\**P* < 0.01, \**P* < 0.05.

As there is a strong correlation between female body size and reproductive output, we first tested for any difference in standard length across selection treatments but found no such effect (up-selected vs. down-selected: ß = 0.0070 [−0.019; 0.036], *P_MCMC_* = 0.50). Number of offspring produced during the first parturition was thereafter compared across selection treatments to assess for potential reproductive costs implicated in the selection for a larger telencephalon. There was no difference in the number of offspring produced across selection treatments (up-selected vs. down-selected: ß = 0.19 [−0.21; 0.59], *P_MCMC_* = 0.28; Figure 3).

**Figure 3.**
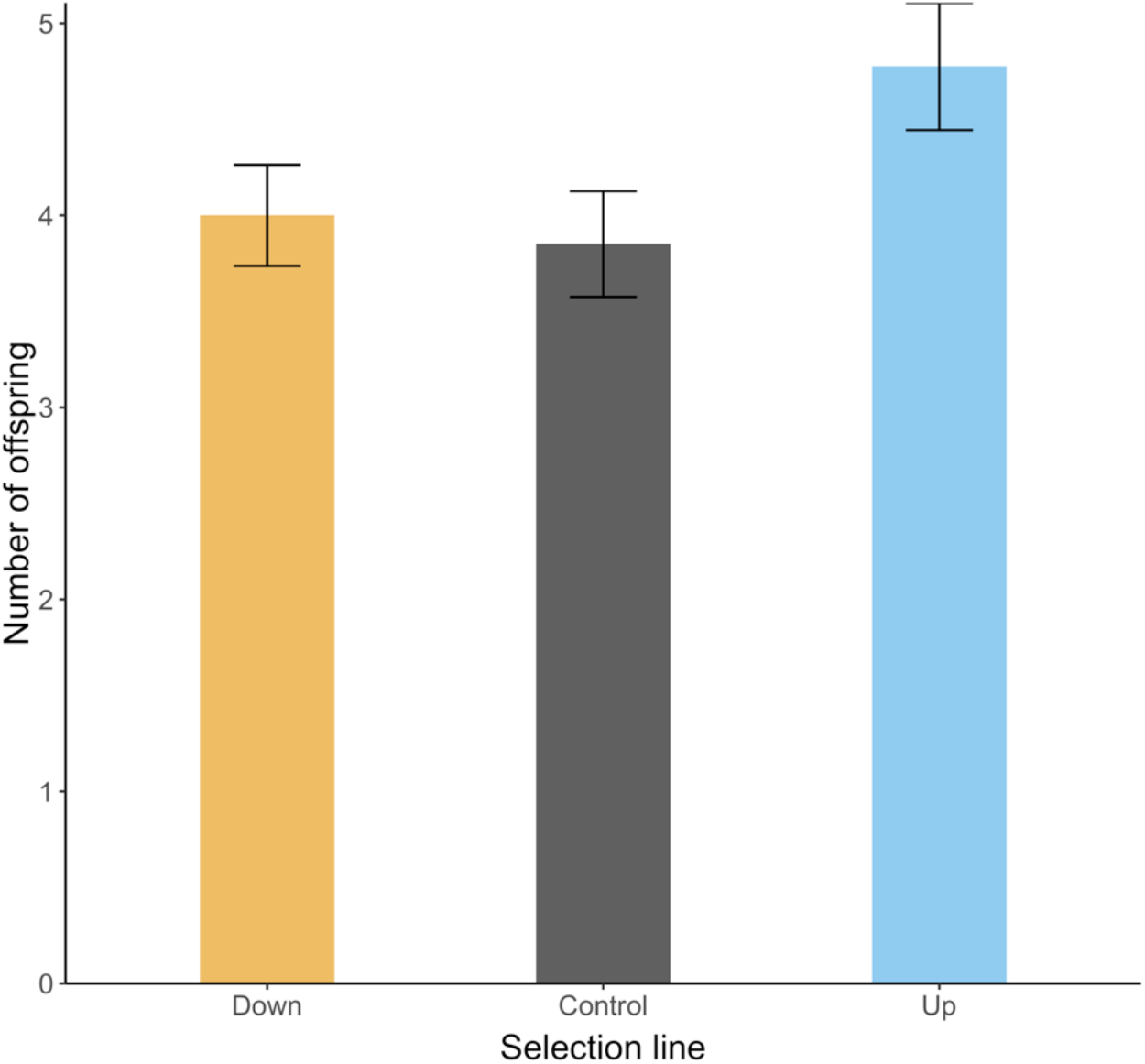
Reproductive output, as assessed by the number of juveniles produced during the first parturition, over the different selection lines. Error bars represent standard error of the mean (sem). No significant differences were detected in generation F4 (p > 0.2).

## Discussion

Our results show that the relative size of a brain region can evolve independently and quickly under strong directional selection. As such, our results provide experimental support for the mosaic brain evolution hypothesis^4,5^. This means that brain regions can display rapid and largely independent evolutionary changes, which further imply that selection can alter individual brain regions in response to specific cognitive demands. Importantly, there is also a contrasting hypothesis to consider, the concerted brain hypothesis, which states that brain regions should evolve in a coordinated manner given developmental constraints^2^. Our results speak against such developmental constraints, at least under strong directional selection.

Comparative studies often find support for a combination of mosaic and concerted evolution^58,59,60^. How do such patterns go together with our experimental results? Artificial selection experiments reveal *what is possible* under strong directional selection, and do not necessarily reflect exactly the patterns and processes that are occurring in wild populations. Hence, while our experiment shows that telencephalon size can evolve highly independently, many opposing selection pressures such as brain cavity space constraints^45^ and/or developmental and energetic constraints^2,46,47^ may impede the signatures of selection for mosaic brain evolution in wild populations. Selection on cognitive ability in wild populations may also target other behavioral and morphological aspects than brain region size^48^. One possible reason why we found it harder to decrease, rather than increase, the size of the telencephalon could be limitations due to cognitive thresholds, whereby a minimum size requirement of the telencephalon is necessary. As mentioned earlier, the teleost telencephalon has known implications in a wide range of cognitive processes, including but not limited to, spatial learning and fear conditioning response. It is therefore conceivable that a fundamental size is required for normal daily functioning, hence curbing the extent to which down-selection can progress. This is in accordance with Haller’s rule, whereby smaller animals tend to possess relatively larger brains due to the minimum absolute brain size required for performing basic cognitive functions^49^.

Realized heritabilities in relative telencephalon size were substantial at least in the up-selected treatments in both sexes but showed high variation. This implies that changes in telencephalon size rests on an important heritable genetic background. While heritability in overall brain volume has been found to be high across species (e.g. humans^50,51^, rhesus monkeys^52^, baboons^53^), genetic influences on brain region sizes have been shown to vary considerably^53–56^. In the previous artificial selection on brain size in guppies, realized heritability of relative brain size was found to be relatively high^19^, suggesting that overall brain size is under strong genetic control. This is congruent with results from human studies where brain volume is significantly influenced by genetic factors, while heritability of individual brain components may vary throughout the lifespan^50^. The measure of heritability used here estimates the proportion of total variance that can be attributed to genetic effects^57^. The fact that we found evidence for considerably higher heritability estimates in the up-selected lines could once again be related to the aforementioned cognitive threshold theory, whereby the requirement for a minimum telencephalon size outweighs genetic influences. Although substantial, the values of realized heritability presented here should be interpreted with some caution. For instance, that the values of realized heritability sometimes lay outside of the normal range of 0-1 may implicate effects of phenotypic plasticity. Given that both brain size and brain region sizes are highly plastic, it is possible that environmental effects may result in a range of region sizes outside of the norm. In addition, it is worth noting that measurement error is generally larger for brain regions than for instance for overall brain size.

We found no evidence for strong associations in the size between different brain regions. Only the region targeted by our selection regime, the telencephalon, changed in size during the experiment. This matches a previous study examining phenotypic and genotypic correlations between brain regions in three-spined stickleback (*Gasterosteus aculeatus*), where Noreikiene, et al. ^58^ found that brain regions shared relatively low genetic correlations. Moreover, in a large-scale quantitative genetic analysis study comprising approximately 10 000 mice, different brain regions were found to be regulated by distinct non-overlapping loci, allowing for the selection of individual brain components^59^. The lack of a link between telencephalon size and other brain regions also speaks against prominent trade-offs between investments into separate regions. This is somewhat surprising, given that neural tissue is one of the most energetically costly tissues^19,29,60,61^. Our experimental design, which targeted relative telencephalon size in relation to the brain remainder, should also have been effective in revealing potential developmental trade-offs with other brain regions. Again, the lack of evidence for any trade-off in this context may stem from high independence and the low genetic correlations between the size of different brain regions. Note that our animals were kept under benign laboratory conditions, which may have masked potential energetic trade-offs. But to fully understand the intricate evolutionary associations between different brain regions will require more work (e.g. rearing of individuals under different food restriction treatments to mimic natural conditions), ideally also into the interconnectivity between regions^17^. Interestingly, the increased telencephalon size in the up-selected lines led to an increase in relative brain size, but only in males (the trend, albeit not statistically significant, was in the same direction also in females (Fig. S2, Table S2)). Substantial changes in the size of specific regions must of course affect also overall brain size unless there are trade-offs with other regions^12^. But comparative analyses have demonstrated that changes in some regions, especially neocortex size and cerebellum size, are linked to changes in brain size (e.g. ^3,12,46^). Although the effect was only evident in males after four generations of selection, our results provide support for a similar pattern also at the intraspecific level. While more work is clearly needed to unveil the developmental pathways and functional implications of the link between telencephalon size and brain size (e.g. ^62^), we propose that mosaic brain evolution can be an important first step also towards larger brain size.

We did not find any evidence for any reproductive trade-off with telencephalon size. This is in contrast to the previous artificial selection experiment on relative brain size in guppies, which found a clear reduction in fecundity (i.e. lower offspring number) in the large-brain lines^19^. Together, these results speak for that mosaic brain evolution of key-regions associated to cognitive demands may be an energy-effective and thus potentially favored way to reduce life history costs during cognitive evolution. As mentioned previously, the comparative and experimental evidence for the costs associated with increased brain size is substantial. However, little is known about the potential costs associated with variation in brain region size. A potential additional route for future research is therefore to investigate if life-history costs are associated also with vertebrate brain region variation in comparative analyses.

Lastly, our comparison of the telencephalon selection lines in relation to 16 recently sampled wild populations of guppies shows that the control-lines are most similar to the wild populations, while up-selected and down-selected lines have larger and smaller telencephali than the wild populations respectively. At the same time, the range of relative telencephalon size was similar in the wild populations as in the selection lines. This result is consistent with that ample genetic variation still exists in the wild-type laboratory strain of guppies used as the basis for this selection experiment^63,64^. Together, these observations suggest that the selection procedure used here has mainly acted within the natural range of telencephalon size variation. Further selection for additional generations will be undertaken to test if artificial selection on a separate brain region can effectively shift its phenotype outside of naturally occurring levels, like what has been done for other morphological traits on numerous occasions^65–68^.

In conclusion, our results show that brain morphology can evolve rapidly in a highly independent fashion under strong directional selection. Our study thus provides experimental support for the mosaic brain evolution hypothesis at the intraspecific level and we propose that mosaic evolution can be an important facilitator of cognitive evolution at the intraspecific level.

## Materials and methods

### Artificial selection for relative telencephalon size

We selected on telencephalon size relative to the size of the rest of the brain, because the alternative, to select on relative telencephalon size against body size, was likely to create larger and smaller relative brain size due to the strong positive correlation between telencephalon size and brain size in vertebrates^69^. Guppies (*Poecilia reticulata*) originating from a high-predation population in Trinidad, and subsequently kept in large populations in the lab since 1998 were used as breeding stock for the starting population. The fish that formed the base populations for these selection lines were housed in large mixed sex groups in three 200 l aquaria. One hundred and twenty breeding pairs were netted out and housed in 3 l aquaria with light-coloured gravel, constant aeration, freely-floating live or plastic plants and > 4 water snails (*Planorbis sp.*) to consume organic waste. The laboratory was maintained at 25 ± 2°C with 12 h light, 12 h dark schedule. Fish were fed six days a week with a diet of flake food and freshly hatched brine shrimp. Given the cannibalistic nature of female guppies, a plastic net with mesh measuring 0.4 × 0.4 cm was placed at the front of the tank, functioning as a safe zone for fry while restricting the access of adult females. Tanks were scanned once every other day for any newborn offspring, which were removed and placed in similar tanks and kept to a maximum of 6 individuals per tank. Among these offspring, males were separated from females at first sign of sexual maturity (i.e. visual detection of gonopodium development) and housed in groups of three to four individuals in similar 3 l aquaria.

At 140 ± 7 (mean ± sd) days of age, 450 of these fish were used to establish the starting population (F_0_) of three independent experimental replicates of 75 breeding pairs each, in similar tanks as described above. Since quantification of brain morphology could not be performed in live fish, we allowed the pairs to produce at least two clutches or at least 12 juveniles per pair, prior to being sacrificed for brain morphology quantification. Breeding pairs were euthanized in a water bath containing an overdose of benzocaine (0.4 g/L), and fixed with 4 % formalin in buffered phosphate buffer saline (PBS) for seven days. The samples were then washed twice in PBS and stored in 4°C awaiting dissection.

Standard length of all fish was recorded to the nearest 0.01 mm, measured from the tip of the snout to the end of the caudal peduncle using digital callipers. Whole brains were dissected out from each fish under a dissecting microscope (Leica MZFLIII), and each individual brain was photographed from four separate views with a digital camera (Leica DFC 490). Based on these separate images, the length, width and height of six major brain regions (i.e. telencephalon, optic tectum, cerebellum, dorsal medulla, hypothalamus and olfactory bulbs) were measured using imageJ software^70^. The volume of each of these brain regions were then determined using the ellipsoid model^71,72^:

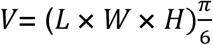

Correspondingly, total brain volume was derived as the sum of the six measured brain regions. To select for relative telencephalon size, we first extracted the residuals from sex-specific regressions of telencephalon volume on the volume of the brain remainder excluding the telencephalon and standardized these to a mean of zero and a standard deviation of one. Based on the sum of the standardized residuals of each pair, offspring from the top and bottom 15 pairs (i.e. top and bottom 20 %) were chosen to form the next generation (F_1_) of up-selected and down-selected lines, respectively across three replicates. Additional offspring from 15 random pairs (F_0_) were likewise used to form the control-lines (see Figure S1 for a detailed schematic outline of the selection procedure). The procedure was largely adapted from a previously successful artificial selection experiment for relative brain size in guppies^19^. Each experimental replicate and selection regime combination was treated independently, thus giving a total of nine distinct populations (three replicates of up-selected, down-selected and control-lines respectively). Within each of the selection lines for each replicate, two females and two males from each of the selected F_0_ pairs were used to form 30 breeding pairs for the next generation (F_1_), resulting in a total of 270 pairs. Again, the top and bottom 15 pairs in terms of relative telencephalon size were chosen for up-selected and down-selected lines while the control-lines were paired randomly in each of the following generations. To prevent inbreeding, breeding pairs never consisted of full siblings.

Similar to the previous generation, F_1_ breeding pairs were allowed to produce at least two clutches or at least 12 juveniles prior to whole brains being dissected out and measured. The selection proceeded in an identical manner as generation F_0_, first by ranking the parents according to the sum of parental residuals and using the respective offspring to propagate the subsequent generations (F_2_, F_3_ and F_4_).

### Brain morphology of wild fish populations

In order to compare the brain morphology of the telencephalon selection lines to wild fish, the brain morphology of wild indigenous female guppies (n = 187) that were obtained from 16 different sites in Trinidad^30^ were quantified in an identical manner as described above. These measurements could thereafter be used as a reference to assess the rate of brain evolution in our selection lines in relation to wild populations.

### Reproductive costs

The development and maintenance of neural tissue incurs high energetic costs^61,73^. In order to examine possible reproductive trade-offs involved in evolving a larger telencephalon, we assessed the possible costs in terms of reproductive output across selection lines. For this, we recorded the total number of juveniles and broods produced by all breeding pairs as a measure of reproductive success.

### Statistical analysis

All statistical analyses were conducted in R statistical software v 3.5.1^74^. We adopted a Bayesian approach in the R package MCMCglmm^75^ to analyze for selection response/effects across lines, with flat priors for the fixed effects and locally uninformative priors for the random effects. We ran each model with 1.65 × 10^6^ iterations, with a thinning interval of 1500 and a burnin of 1.5 × 10^5^, resulting in an effective sample size of at least 1000. All autocorrelations were within the range of −0.1 and 0.1, and were hence deemed acceptable. To avoid statistical confounds resulting from the inclusion of the region being analyzed, we fitted telencephalon volume (log transformed) as the response variable, against the brain remainder excluding the telencephalon (also log transformed) as the covariate. Our model consisted of the following fixed effects: selection treatment (down-selected, control-line, up-selected), brain remainder and sex, with two-way interactions between brain remainder and sex, and between sex and selection treatment. Replicate and the interaction between selection treatment and replicate were included as random factors. The intercept term was set to the regression of telencephalon size on brain remainder for females in the down-selected line.

To test whether selection results stemmed from an indirect selection on overall brain size, we fitted a similar MCMCglmm model with standard length, sex, selection treatment and the interaction between standard length and sex as fixed effects. Random factors included replicate and the interaction between selection treatment and replicate. Analogous models were used to examine for potential correlated selection between the size of telencephalon and other brain regions in each of the five other unselected regions, i.e. optic tectum, cerebellum, dorsal medulla, hypothalamus and olfactory bulbs.

Realized heritability was estimated as the ratio between the mean response to selection and the mean selection differential for each generation, for “up” and “down” selected lines (in comparison to the control-lines), and males and females separately. We ran separate models for each generation and sex combination to estimate the response to selection (calculated as the difference between the “up” or “down” lines as compared to the control-lines). For the models estimating the selection differential from the means of the selected individuals and the population mean (the mean of the guppies in the selection regimes for either smaller or larger telencephalon), we ran models specific to each generation and sex combination. The models estimating the response to selection contained the explanatory variables selection (“up” or “down” and “control”), log brain remainder (total brain volume minus volume of the telencephalon), with replicate and interaction between selection treatment and replicate as random effects. The models estimating the means of the selected individuals contained the explanatory variables selected (“yes”, “no”), log brain remainder as a covariate and replicate line as random effect. To evaluate main effects independently of the covariate, log brain remainder was standardized to a mean of 0 and a standard deviation of 1. All models were fit using MCMCglmm as previously described. The full posterior distribution was included in all analysis.

Trade-offs, in terms of reproductive output, were analyzed by comparing the number of offspring produced during the first parturition for generation F_4_. Since female body size is known to be correlated with fecundity^76^, standard length was included as a covariate in the model. Given the distribution of our response variable (i.e. number of offspring), we fitted a Bayesian model with a Poisson distribution. Hence, our model consisted of the following fixed effects: standard length, selection treatment, and the interaction between the two. Random factors included replicate and the interaction between selection treatment and replicate.

### Ethics

The experiment was performed in accordance with ethical applications approved by the Stockholm Animal Research Ethical Permit Board (Dnr: 223/15, N8/17 and 17362-2019).

## Acknowledgements

We acknowledge valuable feedback from Regina Vega Trejo and David Mitchell, and thank Annika Boussard, Maddi Garate-Olaizola and Vivien Holub for help with animal caretaking. This project was funded by grants to N.K. from the Swedish Research Council (grant 2016-03435), and Knut and Alice Wallenberg Foundation (grant 102 2013.0072).

## Author contributions

NK conceived the idea to the project, SF, AK and NK designed the selection protocol; SF performed the selection experiment; SF, MA, BR and WvdB carried out the data analyses; SB and MA helped with fish maintenance and lab coordination; all authors contributed to the writing of the manuscript.

## Competing interest

The authors declare no competing interests.

## Notes

### Competing Interest Statement

The authors have declared no competing interest.

## References

1 Northcutt, R. G. Understanding vertebrate brain evolution. Integrative and Comparative Biology 42, 743–756, doi:10.1093/icb/42.4.743 (2002).

2 Finlay, B. L. & Darlington, R. B. Linked regularities in the development and evolution of mammalian brains. Science 268, 1578, doi:10.1126/science.7777856 (1995).

3 Striedter, G. F. & Northcutt, R. G. Brains through time: a natural history of vertebrates. (Oxford University Press, 2019).

4 Barton, R. A. & Harvey, P. H. Mosaic evolution of brain structure in mammals. Nature 405, 1055, doi:10.1038/35016580 (2000).

5 Striedter, G. F. Principles of brain evolution. Sinauer Associates (2005).

6 de Winter, W. & Oxnard, C. E. Evolutionary radiations and convergences in the structural organization of mammalian brains. Nature 409, 710–714, doi:10.1038/35055547 (2001).

7 Reader, S. M. & Laland, K. N. Social intelligence, innovation, and enhanced brain size in primates. Proceedings of the National Academy of Sciences 99, 4436 (2002).

8 Lefebvre, L., Reader, S. M. & Sol, D. Brains, innovations and evolution in birds and primates. Brain, Behavior and Evolution 63, 233–246, doi:10.1159/000076784 (2004).

9 Safi, K. & Dechmann, D. K. Adaptation of brain regions to habitat complexity: a comparative analysis in bats (Chiroptera). Proceedings of the Royal Society B: Biological Sciences 272, 179 (2005).

10 Gonzalez-Voyer, A., Winberg, S. & Kolm, N. Brain structure evolution in a basal vertebrate clade: evidence from phylogenetic comparative analysis of cichlid fishes. BMC Evolutionary Biology 9, 238, doi:10.1186/1471-2148-9-238 (2009).

11 Gómez-Robles, A., Hopkins, W. D. & Sherwood, C. C. Modular structure facilitates mosaic evolution of the brain in chimpanzees and humans. Nature Communications 5, 4469, doi:10.1038/ncomms5469 (2014).

12 Sukhum, K. V., Shen, J. & Carlson, B. A. Extreme enlargement of the cerebellum in a clade of teleost fishes that evolved a novel active sensory system. Current Biology 28, 3857–3863.e3853, doi:https://doi.org/10.1016/j.cub.2018.10.038 (2018).

13 Krebs, J. R., Sherry, D. F., Healy, S. D., Perry, V. H. & Vaccarino, A. L. Hippocampal specialization of food-storing birds. Proceedings of the National Academy of Sciences 86, 1388, doi:10.1073/pnas.86.4.1388 (1989).

14 Sherry, D. F., Vaccarino, A. L., Buckenham, K. & Herz, R. S. The hippocampal complex of food-storing birds. Brain, behavior and evolution 34, 308–317 (1989).

15 Garamszegi, L. Z. & Eens, M. The evolution of hippocampus volume and brain size in relation to food hoarding in birds. Ecology Letters 7, 1216–1224, doi:10.1111/j.1461-0248.2004.00685.x (2004).

16 Garamszegi, L. Z. & Eens, M. Brain space for a learned task: strong intraspecific evidence for neural correlates of singing behavior in songbirds. Brain Research Reviews 44, 187–193, doi:https://doi.org/10.1016/j.brainresrev.2003.12.001 (2004).

17 Iwaniuk, A. N., Dean, K. M. & Nelson, J. E. A mosaic pattern characterizes the evolution of the avian brain. Proceedings of the Royal Society of London. Series B: Biological Sciences 271, S148–S151, doi:10.1098/rsbl.2003.0127 (2004).

18 Markina, N. V., Popova, N. V. & Poletaeva, II. Interstrain differences in the the behavior of mice selected for greater and lesser brain mass. Zhurnal vysshei nervnoi deiatelnosti imeni I P Pavlova 49, 59–67 (1999).

19 Kotrschal, A. et al. Artificial selection on relative brain size in the guppy reveals costs and benefits of evolving a larger brain. Current biology : CB 23, 168–171, doi:10.1016/j.cub.2012.11.058 (2013).

20 Kotrschal, A., Corral-Lopez, A., Amcoff, M. & Kolm, N. A larger brain confers a benefit in a spatial mate search learning task in male guppies. Behavioral Ecology 26, 527–532 (2015).

21 Buechel, S. D., Boussard, A., Kotrschal, A., van der Bijl, W. & Kolm, N. Brain size affects performance in a reversal-learning test. Proceedings of the Royal Society B: Biological Sciences 285(2018).

22 Perepelkina, O. V., Lilp, I. G., Tarasova, A. Y., Golibrodo, V. A. & Poletaeva, I. I. Changes in cognitive abilities of laboratory mice as a result of artificial selection. The Russian Journal of Cognitive Science 2, 29–35 (2015).

23 Perepelkina, O. V., Tarasova, A. Y., Ogienko, N. A., Lil’p, I. G. & Poletaeva, I. I. Brain weight and cognitive abilities of laboratory mice. Biology Bulletin Reviews 10, 91–101, doi:10.1134/S2079086420020061 (2020).

24 Corral-López, A. et al. Female brain size affects the assessment of male attractiveness during mate choice. Science Advances 3(2017).

25 Corral-López, A., Romensky, M., Kotrschal, A., Buechel, S. D. & Kolm, N. Brain size affects responsiveness in mating behaviour to variation in predation pressure and sex ratio. Journal of Evolutionary Biology 33, 165–177, doi:10.1111/jeb.13556 (2020).

26 Kotrschal, A. et al. Brain size affects female but not male survival under predation threat. Ecology Letters 18, 646–652, doi:10.1111/ele.12441 (2015).

27 van der Bijl, W., Thyselius, M., Kotrschal, A. & Kolm, N. Brain size affects the behavioural response to predators in female guppies (*Poecilia reticulata*). Proceedings of the Royal Society B: Biological Sciences 282(2015).

28 Kotrschal, A., Kolm, N. & Penn, D. J. Selection for brain size impairs innate, but not adaptive immune responses. Proceedings of the Royal Society B: Biological Sciences 283(2016).

29 Kotrschal, A., Corral-Lopez, A. & Kolm, N. Large brains, short life: selection on brain size impacts intrinsic lifespan. Biology Letters 15, 20190137, doi:10.1098/rsbl.2019.0137 (2019).

30 Kotrschal, A., Deacon, A. E., Magurran, A. E. & Kolm, N. Predation pressure shapes brain anatomy in the wild. Evolutionary Ecology 31, 619–633, doi:10.1007/s10682-017-9901-8 (2017).

31 Huber, R., van Staaden, M. J., Kaufman, L. S. & Liem, K. F. Microhabitat use, trophic patterns, and the evolution of brain structure in African Cichlids. Brain, Behavior and Evolution 50, 167–182, doi:10.1159/000113330 (1997).

32 Aiello, L. C. & Wheeler, P. The expensive-tissue hypothesis: the brain and the digestive System in human and primate evolution. Current Anthropology 36, 199–221, doi:10.1086/204350 (1995).

33 Logan, C. J. et al. Beyond brain size: Uncovering the neural correlates of behavioral and cognitive specialization. Comparative Cognition & Behavior Reviews 13, 55–89, doi:10.3819/CCBR.2018.130008 (2018).

34 Portavella, M., Vargas, J. P., Torres, B. & Salas, C. The effects of telencephalic pallial lesions on spatial, temporal, and emotional learning in goldfish. Brain Research Bulletin 57, 397–399, doi:https://doi.org/10.1016/S0361-9230(01)00699-2 (2002).

35 Broglio, C., Rodríguez, F. & Salas, C. Spatial cognition and its neural basis in teleost fishes. Fish and Fisheries 4, 247–255, doi:10.1046/j.1467-2979.2003.00128.x (2003).

36 Broglio, C. et al. Hallmarks of a common forebrain vertebrate plan: specialized pallial areas for spatial, temporal and emotional memory in actinopterygian fish. Brain Research Bulletin 66, 277–281, doi:https://doi.org/10.1016/j.brainresbull.2005.03.021 (2005).

37 Corbin, J. G., Nery, S. & Fishell, G. Telencephalic cells take a tangent: non-radial migration in the mammalian forebrain. Nature Neuroscience 4, 1177–1182, doi:10.1038/nn749 (2001).

38 Guillemot, F. Cellular and molecular control of neurogenesis in the mammalian telencephalon. Current Opinion in Cell Biology 17, 639–647, doi:https://doi.org/10.1016/j.ceb.2005.09.006 (2005).

39 Houde, A. Sex, color, and mate choice in guppies. . Princeton University Press. (1997).

40 Reznick, D. N., Butler IV, M. J. & Rodd, H. Life-history evolution in guppies. VII. The comparative ecology of high- and low-predation environments. The American Naturalist 157, 126–140, doi:10.1086/318627 (2001).

41 Reznick, D. N. Life history evolution in guppies (*Poecilia reticulata*): guppies as a model for studying the evolutionary biology of aging. Experimental gerontology 32, 245–258, doi:10.1016/S0531-5565(96)00129-5 (1997).

42 Harris, S., Ramnarine, I. W., Smith, H. G. & Pettersson, L. B. Picking personalities apart: estimating the influence of predation, sex and body size on boldness in the guppy *Poecilia reticulata*. Oikos 119, 1711–1718, doi:10.1111/j.1600-0706.2010.18028.x (2010).

43 Isler, K. & van Schaik, C. P. The expensive brain: a framework for explaining evolutionary changes in brain size. Journal of Human Evolution 57, 392–400, doi:https://doi.org/10.1016/j.jhevol.2009.04.009 (2009).

44 Burns, J. G., Saravanan, A. & Helen Rodd, F. Rearing environment affects the brain size of guppies: lab-reared guppies have smaller brains than wild-caught guppies. Ethology 115, 122–133, doi:doi:10.1111/j.1439-0310.2008.01585.x (2009).

45 Striedter, G. F. & Northcutt, R. G. Head size constrains forebrain development and evolution in ray-finned fishes. Evolution & Development 8, 215–222, doi:10.1111/j.1525-142X.2006.00091.x (2006).

46 Finlay, B. L., Darlington, R. B. & Nicastro, N. Developmental structure in brain evolution. Behavioral and Brain Sciences 24, 263–278, doi:10.1017/S0140525X01003958 (2001).

47 Niven, J. E. & Laughlin, S. B. Energy limitation as a selective pressure on the evolution of sensory systems. Journal of Experimental Biology 211, 1792, doi:10.1242/jeb.017574 (2008).

48 Thornton, A. & Lukas, D. Individual variation in cognitive performance: developmental and evolutionary perspectives. Philosophical Transactions of the Royal Society B: Biological Sciences 367, 2773–2783, doi:10.1098/rstb.2012.0214 (2012).

49 Eberhard, W. G. & Wcislo, W. T. in Advances in Insect Physiology Vol. 40 (ed Jérôme Casas) 155–214 (Academic Press, 2011).

50 Batouli, S. A. H., Trollor, J. N., Wen, W. & Sachdev, P. S. The heritability of volumes of brain structures and its relationship to age: a review of twin and family studies. Ageing Research Reviews 13, 1–9, doi:https://doi.org/10.1016/j.arr.2013.10.003 (2014).

51 Bartley, A. J., Jones, D. W. & Weinberger, D. R. Genetic variability of human brain size and cortical gyral patterns. Brain 120, 257–269, doi:10.1093/brain/120.2.257 (1997).

52 Cheverud, J. M. et al. Heritability of brain size and surface features in rhesus macaques (*Macaca mulatta*). Journal of Heredity 81, 51–57, doi:10.1093/oxfordjournals.jhered.a110924 (1990).

53 Rogers, J. et al. Heritability of brain volume, surface area and shape: an MRI study in an extended pedigree of baboons. Hum Brain Mapp 28, 576–583, doi:10.1002/hbm.20407 (2007).

54 Peper, J. S., Brouwer, R. M., Boomsma, D. I., Kahn, R. S. & Hulshoff Pol, H. E. Genetic influences on human brain structure: a review of brain imaging studies in twins. Human Brain Mapping 28, 464–473, doi:10.1002/hbm.20398 (2007).

55 Sullivan, E. V., Pfefferbaum, A., Swan, G. E. & Carmelli, D. Heritability of hippocampal size in elderly twin men: equivalent influence from genes and environment. Hippocampus 11, 754–762, doi:10.1002/hipo.1091 (2001).

56 Airey, D. C., Castillo-Juarez, H., Casella, G., Pollak, E. J. & DeVoogd, T. J. Variation in the volume of zebra finch song control nuclei is heritable: developmental and evolutionary implications. Proceedings of the Royal Society of London. Series B: Biological Sciences 267, 2099–2104, doi:10.1098/rspb.2000.1255 (2000).

57 Lynch, M. & Walsh, B. Genetics and analysis of quantitative traits. Vol. 1 (Sinauer Sunderland, MA, 1998).

58 Noreikiene, K. et al. Quantitative genetic analysis of brain size variation in sticklebacks: support for the mosaic model of brain evolution. Proceedings of the Royal Society B: Biological Sciences 282, 20151008, doi:10.1098/rspb.2015.1008 (2015).

59 Hager, R., Lu, L., Rosen, G. D. & Williams, R. W. Genetic architecture supports mosaic brain evolution and independent brain–body size regulation. Nature Communications 3, 1079, doi:10.1038/ncomms2086 (2012).

60 Kotrschal, A., Kolm, N. & Penn, D. J. Selection for brain size impairs innate, but not adaptive immune responses. Proceedings of the Royal Society B: Biological Sciences 283, 20152857, doi:10.1098/rspb.2015.2857 (2016).

61 Laughlin, S. B., de Ruyter van Steveninck, R. R. & Anderson, J. C. The metabolic cost of neural information. Nature Neuroscience 1, 36–41, doi:10.1038/236 (1998).

62 Charvet, C. J., Striedter, G. F. & Finlay, B. L. Evo-devo and brain scaling: candidate developmental mechanisms for variation and constancy in vertebrate brain evolution. Brain, Behavior and Evolution 78, 248–257, doi:10.1159/000329851 (2011).

63 Robertson, A. Inbreeding in artificial selection programmes. Genetical Research 2, 189–194, doi:10.1017/S0016672300000690 (1961).

64 Chen, N., Luo, X., Lu, C., Ke, C. & You, W. Effects of artificial selection practices on loss of genetic diversity in the Pacific abalone, *Haliotis discus hannai*. Aquaculture Research 48, 4923–4933, doi:10.1111/are.13311 (2017).

65 Beldade, P., Koops, K. & Brakefield, P. M. Developmental constraints versus flexibility in morphological evolution. Nature 416, 844–847, doi:10.1038/416844a (2002).

66 Hansen, S. W. Selection for behavioural traits in farm mink. Applied Animal Behaviour Science 49, 137–148, doi:https://doi.org/10.1016/0168-1591(96)01045-3 (1996).

67 von Schantz, T. et al. Artificial selection for increased comb size and its effects on other sexual characters and viability in *Gallus domesticus* (the domestic chicken). Heredity 75, 518–529, doi:10.1038/hdy.1995.168 (1995).

68 Jackson, S. & Diamond, J. Metabolic and digestive responses to artificial selection in chickens. Evolution 50, 1638–1650, doi:10.1111/j.1558-5646.1996.tb03936.x (1996).

69 Yopak, K. E. et al. A conserved pattern of brain scaling from sharks to primates. Proceedings of the National Academy of Sciences 107, 12946, doi:10.1073/pnas.1002195107 (2010).

70 Schindelin, J. et al. Fiji: an open-source platform for biological-image analysis. Nature Methods 9, 676, doi:10.1038/nmeth.2019 (2012).

71 Pollen, A. A. et al. Environmental complexity and social organization sculpt the brain in Lake Tanganyikan cichlid fish. Brain, Behavior and Evolution 70, 21–39 (2007).

72 White, G. E. & Brown, C. Variation in brain morphology of intertidal gobies: a comparison of methodologies used to quantitatively assess brain volumes in fish. Brain, Behavior and Evolution 85, 245–256, doi:10.1159/000398781 (2015).

73 Mink, J. W., Blumenschine, R. J. & Adams, D. B. Ratio of central nervous system to body metabolism in vertebrates: its constancy and functional basis. American Journal of Physiology-Regulatory, Integrative and Comparative Physiology 241, R203–R212, doi:10.1152/ajpregu.1981.241.3.R203 (1981).

74 Team, R. RStudio: Integrated Development for R. RStudio, Inc., Boston, MA (2015).

75 Hadfield, J. D. MCMC Methods for Multi-Response Generalized Linear Mixed Models: The MCMCglmm R Package. Journal of Statistical Software 33, 1–22 (2010).

76 Roff, D. The evolution of life histories: theory and analysis. The evolution of life histories: theory and analysis. (1992).

