## Supplementary Figures and Tables for "Rapid mosaic brain evolution under artificial selection for relative telencephalon size in the guppy (*Poecilia reticulata*)"

**Supplemental inventory**

**Figure S1.** This figure illustrates the artificial selection procedure for relative telencephalon size.

**Figure S2.** This figure shows overall changes in brain morphology in response to directed selection for telencephalon size in female guppies.

**Figure S3**. This figure shows overall changes in brain morphology in response to directed selection for telencephalon size in male guppies.

**Figure S4.** This figure shows the range of relative telencephalon sizes in females across 16 wild Trinidadian populations and generation $F_4$ females.

**Table S1.** This table shows the model output for overall changes in brain morphology in response to selection for relative telencephalon size.

**Table S2.** This table shows the model output for changes in brain size (relative to standard length) in both females and males in response to selection for relative telencephalon size.

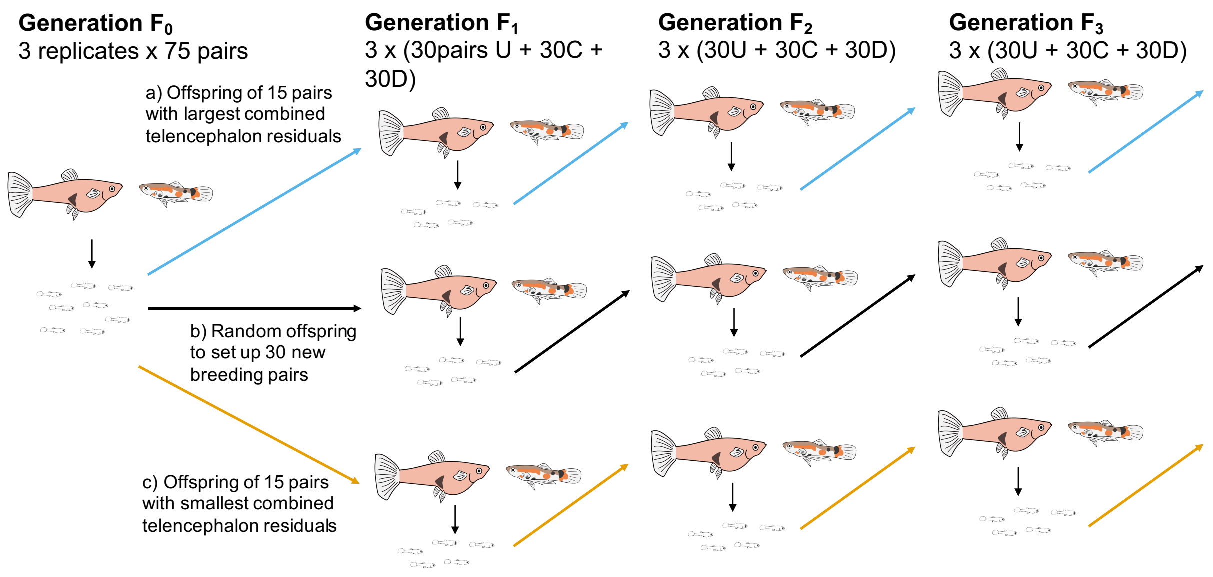


**Figure S1.** Artificial selection procedure for relative telencephalon size. Different selection lines are indicated by the different coloured arrows (i.e. blue for up-selection, orange for control and black for down-selected). Within each generation, 3 independent replicates of 30 breeding pairs for each selection line were set up.

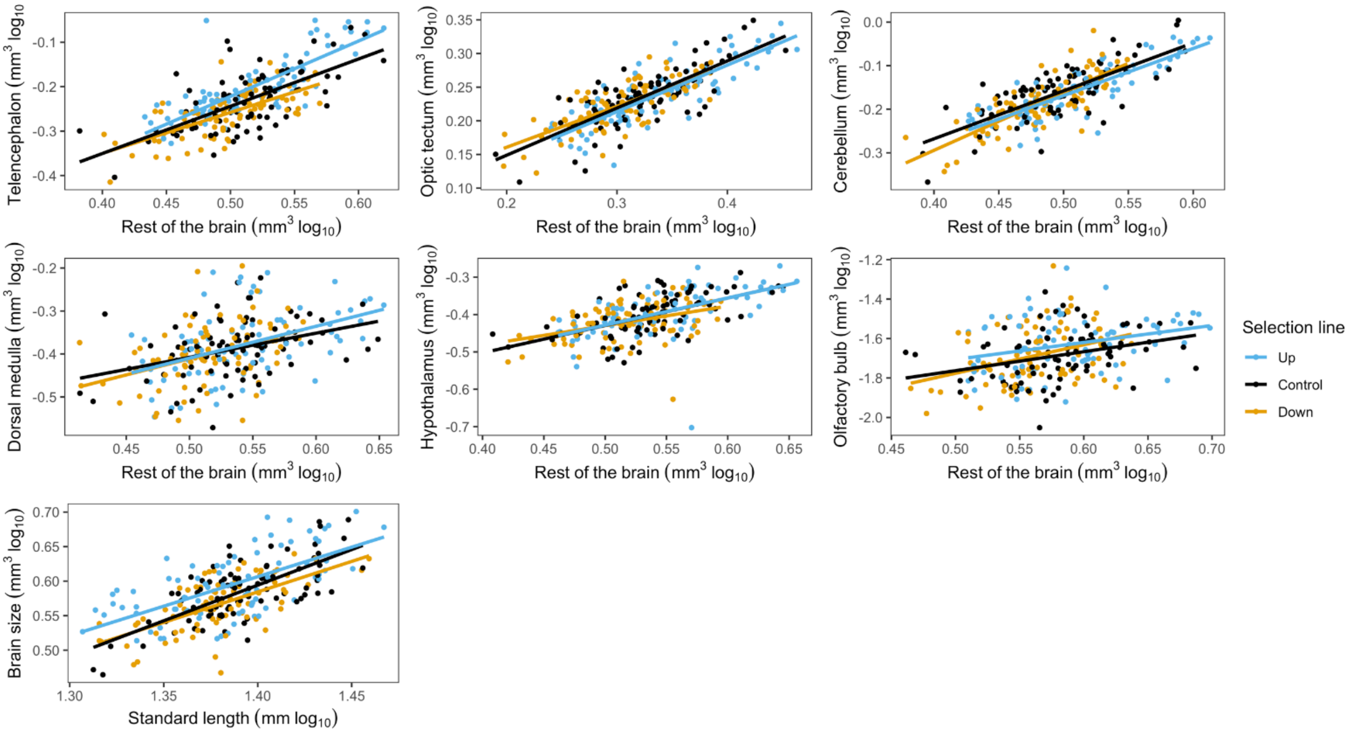


**Figure S2.** Overall changes in brain size and brain sub regions in response to directed selection on relative telencephalon size in generation $F_4$ females. Shown in the figures are the raw data of each brain region (log transformed) on the y-axis and brain remainder (log transformed) on the x-axis.

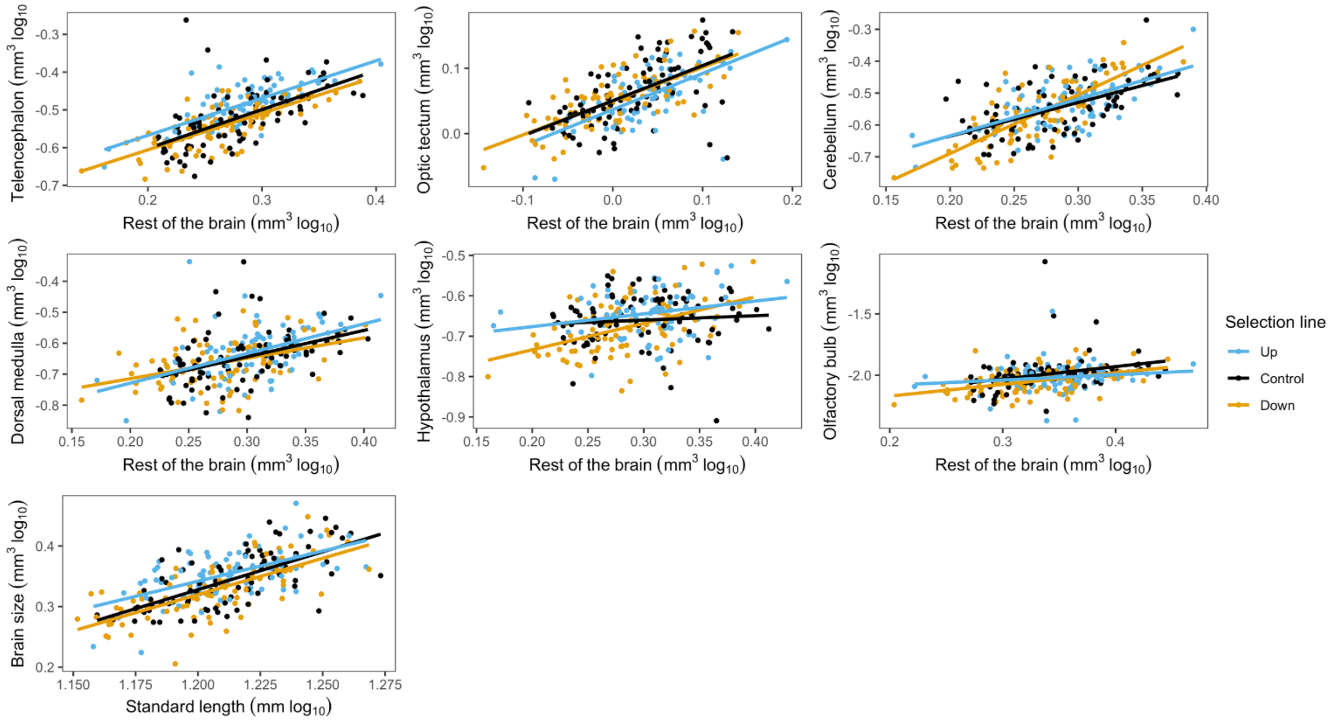


**Figure S3.** Overall changes in brain size and brain sub regions in response to directed selection on relative telencephalon size in generation $F_4$ males. Depicted in the figures are the raw data of each brain region (log transformed) on the y-axis and brain remainder (log transformed) on the x-axis.

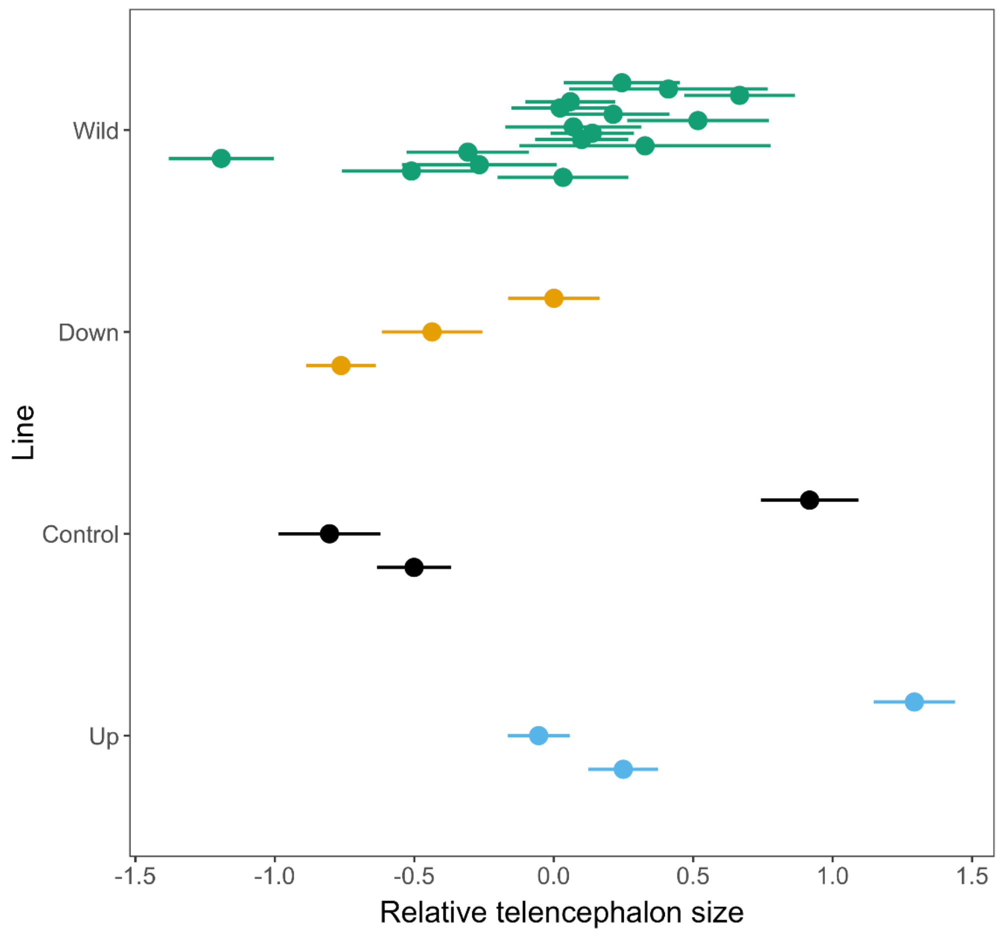


**Figure S4.** Range of relative telencephalon size in females from each selection treatment ($F_4$) and wild Trinidadian populations. Shown are the mean relative telencephalon size and standard error of the mean (sem), obtained from standardized residuals of telencephalon volume regressed on the brain remainder (total brain volume minus telencephalon volume), across lines.

**Table S1.** Model output for overall changes in brain morphology in generation $F_4$ adults in response to selection for relative telencephalon size, presented with mean estimates and 95 % credible intervals (CI). The intercept is set to the intercept of the regression of log sub region size on log brain remainder (brain volume minus volume of region of interest) for the down-selected line in females. The mean values presented in the table represent differences between the intercept (down-selected females) and the specified variable (conforming to the default contrast matrix in R).

| | ESTIMATE (ß) | LOWER CI | UPPER CI | $P_{MCMC}$ |
|---|---|---|---|---|
| **Overall brain** | | | | |
| Intercept | -0.677 | -0.898 | -0.445 | 0.002** |
| Standard length | 0.897 | 0.776 | 1.02 | <0.001*** |
| Sex Male | -0.364 | -0.598 | -0.126 | 0.004** |
| Unselected control | 0.00756 | -0.00914 | 0.0260 | 0.278 |
| Up-selected line | 0.0233 | 0.00730 | 0.0423 | 0.022* |
| Standard length:Sex Male | 0.232 | 0.0357 | 0.408 | 0.016* |
| Sex Male:Unselected control | 0.000502 | -0.0110 | 0.0129 | 0.926 |
| Sex Male:Up-selected line | -0.00136 | -0.0139 | 0.0107 | 0.842 |
| **Optic tectum** | | | | |
| Intercept | 0.00619 | -0.0108 | 0.0701 | 0.138 |
| Rest of brain | 0.656 | 0.575 | 0.737 | <0.001*** |
| Sex | -0.0251 | -0.00217 | 0.0491 | 0.076 |
| Unselected control | -0.00141 | -0.0139 | 0.0162 | 0.846 |
| Up-selected line | -0.00606 | -0.0225 | 0.00969 | 0.370 |

| | | | | |
|---|---|---|---|---|
| Rest of brain:Sex | -0.137 | -0.246 | -0.0256 | 0.010** |
| Sex:Unselected control | 0.00254 | -0.0104 | 0.0168 | 0.676 |
| Sex:Up-selected line | -0.00488 | -0.0191 | 0.00988 | 0.518 |
| **Cerebellum** | | | | |
| Intercept | -0.740 | -0.858 | -0.548 | <0.001*** |
| Rest of brain | 1.13 | 0.970 | 1.29 | <0.001*** |
| Sex | -0.208 | -0.294 | -0.131 | <0.001*** |
| Unselected control | 0.00678 | -0.0625 | 0.0539 | 0.758 |
| Up-selected line | -0.00500 | -0.0625 | 0.0543 | 0.790 |
| Rest of brain:Sex | 0.264 | 0.0756 | 0.492 | 0.010* |
| Sex:Unselected control | -0.0116 | -0.0326 | 0.00984 | 0.292 |
| Sex:Up-selected line | 0.00844 | -0.0118 | 0.0316 | 0.456 |
| **Dorsal medulla** | | | | |
| Intercept | -0.725 | -1.03 | -0.454 | 0.010** |
| Rest of brain | 0.628 | 0.416 | 0.817 | <0.001*** |
| Sex | -0.153 | -0.262 | -0.0368 | 0.014* |
| Unselected control | -0.00234 | -0.0423 | 0.0333 | 0.854 |
| Up-selected line | 0.00582 | -0.0331 | 0.0456 | 0.700 |
| Rest of brain:Sex | 0.143 | -0.118 | 0.404 | 0.312 |
| Sex:Unselected control | 0.00329 | -0.0236 | 0.0324 | 0.842 |
| Sex:Up-selected line | 0.00769 | -0.0217 | 0.0361 | 0.654 |
| **Hypothalamus** | | | | |
| Intercept | -0.714 | -0.834 | -0.610 | 0.008** |

| | | | | |
|---|---|---|---|---|
| Rest of brain | 0.577 | 0.423 | 0.751 | <0.001*** |
| Sex | -0.0413 | -0.132 | 0.0529 | 0.396 |
| Unselected control | 0.00802 | -0.0191 | 0.0395 | 0.542 |
| Up-selected line | 0.0110 | -0.0163 | 0.0408 | 0.442 |
| Rest of brain:Sex | -0.291 | -0.517 | -0.0966 | 0.006** |
| Sex:Unselected control | 0.00740 | -0.0142 | 0.0312 | 0.651 |
| Sex:Up-selected line | 0.0216 | -0.00224 | 0.0449 | 0.074 |
| **Olfactory bulbs** | | | | |
| Intercept | -2.27 | -2.73 | -1.88 | <0.001*** |
| Rest of brain | 1.02 | 0.657 | 1.34 | <0.001*** |
| Sex | -0.0580 | -0.280 | 0.144 | 0.600 |
| Unselected control | -0.0170 | -0.0846 | 0.0504 | 0.514 |
| Up-selected line | 0.0290 | -0.0440 | 0.103 | 0.312 |
| Rest of brain:Sex | -0.202 | -0.668 | 0.227 | 0.386 |
| Sex:Unselected control | 0.0640 | 0.0165 | 0.110 | 0.006** |
| Sex:Up-selected line | -0.0149 | -0.0639 | 0.0307 | 0.542 |

**Table S2.** Sex-specific changes in relative brain size in generation $F_4$ females and males in response to selection for relative telencephalon size, presented with mean estimates and 95 % credible intervals (CI). The intercept is set to the intercept of the regression of log brain size on log body size (standard length) for the down-selected line in both females and males, respectively.

| **Females** | | | | |
|---|---|---|---|---|
| **Fixed terms** | Estimate (ß) | Lower CI | Upper CI | $P_{MCMC}$ |
| Intercept | -0.775 | -1.01 | -0.561 | 0.004 ** |
| Standard length | 0.969 | 0.838 | 1.10 | <0.001 *** |
| Unselected control | 0.00773 | -0.0218 | 0.0461 | 0.550 |
| Up-selected | 0.0251 | -0.00703 | 0.0576 | 0.098 |
| **Random terms** | Variance | Lower CI | Upper CI | *P* |
| Residual | 0.000745 | 0.000617 | 0.000880 | NA |
| Selection treatment:Replicate | 0.000451 | $1.19e^{-5}$ | 0.00168 | NA |
| Replicate | 0.0369 | $5.63e^{-11}$ | 0.0907 | NA |
| **Males** | | | | |
| **Fixed terms** | Estimate (ß) | Lower CI | Upper CI | $P_{MCMC}$ |
| Intercept | -0.978 | -1.20 | -0.756 | 0.002 ** |
| Standard length | 1.08 | 0.927 | 1.25 | <0.001 *** |
| Unselected control | 0.00802 | -0.00980 | 0.0261 | 0.274 |
| Up-selected | 0.0217 | 0.00521 | 0.0405 | 0.028 * |
| **Random terms** | Variance | Lower CI | Upper CI | *P* |
| Residual | 0.000901 | 0.000749 | 0.00106 | NA |

| | | | | |
|---|---|---|---|---|
| Selection treatment:Replicate | 0.000109 | $1.13e^{-10}$ | 0.000411 | NA |
| Replicate | 0.0525 | $3.72e^{-7}$ | 0.0886 | NA |

**Statistical analysis**

Selection response was also analyzed using Restricted Maximum Likelihood, but results were found to concur with those from the Bayesian models. Briefly, to examine the response to selection for relative telencephalon size, we fitted a linear mixed effect model (using the package lme4[53]) with the fixed effects brain remainder as a covariate, selection regime (“down”, “control” and “up”) and sex (“female”, “male”). Random factors included replicate and replicate nested within selection treatment. (lme4 syntax for R model: telencephalon ~ rest of the brain + selection treatment*sex + (1|selection treatment:replicate) + (1|replicate)).

To examine for associated changes in brain morphology in response to directed selection for relative telencephalon size, volumes of each of the remaining five brain sub regions (i.e. optic tectum, cerebellum, dorsal medulla, hypothalamus and olfactory bulbs) were compared across selection lines. An analogous model was fitted for each of the brain sub regions as follows: lmer(brain region ~ rest of the brain + selection treatment*sex + (1|selection treatment:replicate) + (1|replicate)).

In order to generate measures of evolutionary change in relative telencephalon size, we computed standardized residuals of telencephalon volume regressed on the volume of the brain remainder. Residuals were extracted from a model that included both selected (i.e. down-, up-selected and unselected controls) and wild fish, and the mean of each group was determined (R syntax: stdres(telencephalon ~ rest of the brain). Thereafter, we compared the mean telencephalon size as well as the range of data between the selection lines and wild Trinidadian fish to examine how artificial selection altered telencephalon size and telencephalon size range relative to the natural span.
